# Attenuated CaV1.2-BK channel protein interaction in bipolar disorder

**DOI:** 10.1101/2025.08.08.669447

**Authors:** Rupali Srivastava, Nathaniel Genduso, Koko Ishizuka, Akira Sawa

## Abstract

Bipolar disorder (BP) is a severe mental illness featured with potential cycling of depressive and manic episodes. The recurrent cycling is known to be associated with poor prognosis. Thus, it is important to decipher the pathophysiological mechanism that may possibly underlie the cycling. In our study, by using BP patient-derived neurons, we discovered the delay of time-to-peak in depolarization-associated intracellular calcium dynamics. This cellular deficit is selectively associated with the number of manic episodes in BP patients, suggesting that this deficit may participate in the mechanism of clinical cycling. We also observed an attenuated protein interaction between the calcium and voltage-activated potassium (BK) channel and the CaV1.2 L-type VGCC in BP patients. Pharmacological and molecular interventions that target CaV1.2-BK protein interaction, directly to BK channel for the amelioration of this attenuated protein interaction rescues the disease-associated intracellular calcium deficit. To test the involvement of BK channel in cycling-associated behaviors, we generated a novel animal model.

## INTRODUCTION

Bipolar disorder (BP) is a severe mental illness featured with potential cycling of depressive and manic episodes^**1,2**^. Epidemiological data show that BP has a lifetime prevalence of 4-5% in the United States. At least 6 to 7% of BP patients finally die through suicide; there is evidence that suicide rates in BP patients are 20 to 30 times as high as the rates in the general population. The onset of the disease is frequently in young adulthood, leading to chronic disabilities. In particular, the recurrent cycling is known to be associated with poor prognosis. Thus, it is important to decipher the pathophysiological mechanism that may possibly underlie the cycling. The deficits in intracellular calcium [Ca^2+^]_i_ homeostasis have been classically suggested in BP. Recent genome wide association studies have also supported the idea by highlighting the genes coding for L-type voltage gated calcium channels (VGCC)^**3,4**^. Additional discoveries utilizing BP patient-derived neuronal models have supported this pathophysiological hypothesis^**5**^.

Nevertheless, this hypothesis has not be fully developed for drug discovery and translation.

In our study, by using BP patient-derived neurons, we discovered the delay of time-to-peak in depolarization-associated [Ca^2+^]_i_ dynamics. This cellular deficit is selectively associated with the number of manic episodes in BP patients, suggesting that this deficit may participate in the mechanism of clinical cycling. We also observed an attenuated protein interaction between the calcium and voltage-activated potassium (BK) channel and the CaV1.2 L-type VGCC in BP patients. Pharmacological amelioration of this attenuated protein interaction rescues the disease-associated [Ca^2+^]_i_ deficit. To test the involvement of BK channel in cycling-associated behaviors, we generated a novel animal model.

## RESULTS AND DISCUSSION

### Deficit in the intracellular Ca^2+^ homeostasis in BP neurons

Based on our hypothesis that neuronal cells from BP patients may have alternations in the intracellular Ca^2+^ ([Ca^2+^]_i_) homeostasis, we examined induced neuronal cells (iN cells) from 25 sporadic BP patients in comparison with those from 26 healthy subjects (**Fig. 1a**). Although Fura2a did not detect a difference of baseline [Ca^2+^]_i_ between these two groups (data not shown), we observed a delay of [Ca^2+^]_i_ time-to-peak in a high potassium-evoked depolarization (**Fig. 1b, c**). Next, we investigated whether this [Ca^2+^]_i_ deficit is associated with clinical manifestations in BP patients. Notably, the number of manic episodes was the only variable among many clinical factors with a significant correlation with the cellular characteristic, time-to-peak, after correction of multiple comparisons (**Fig. 1d**). This cellular deficit was also reproducibly seen in iPS cell-derived excitatory neurons (**Fig. 1e**). By using multiple pharmacological agents, we underscored that L-type voltage-gated calcium channel (VGCC) plays a key role in this [Ca^2+^]_i_ deficit (data not shown).

**Fig. 1:**
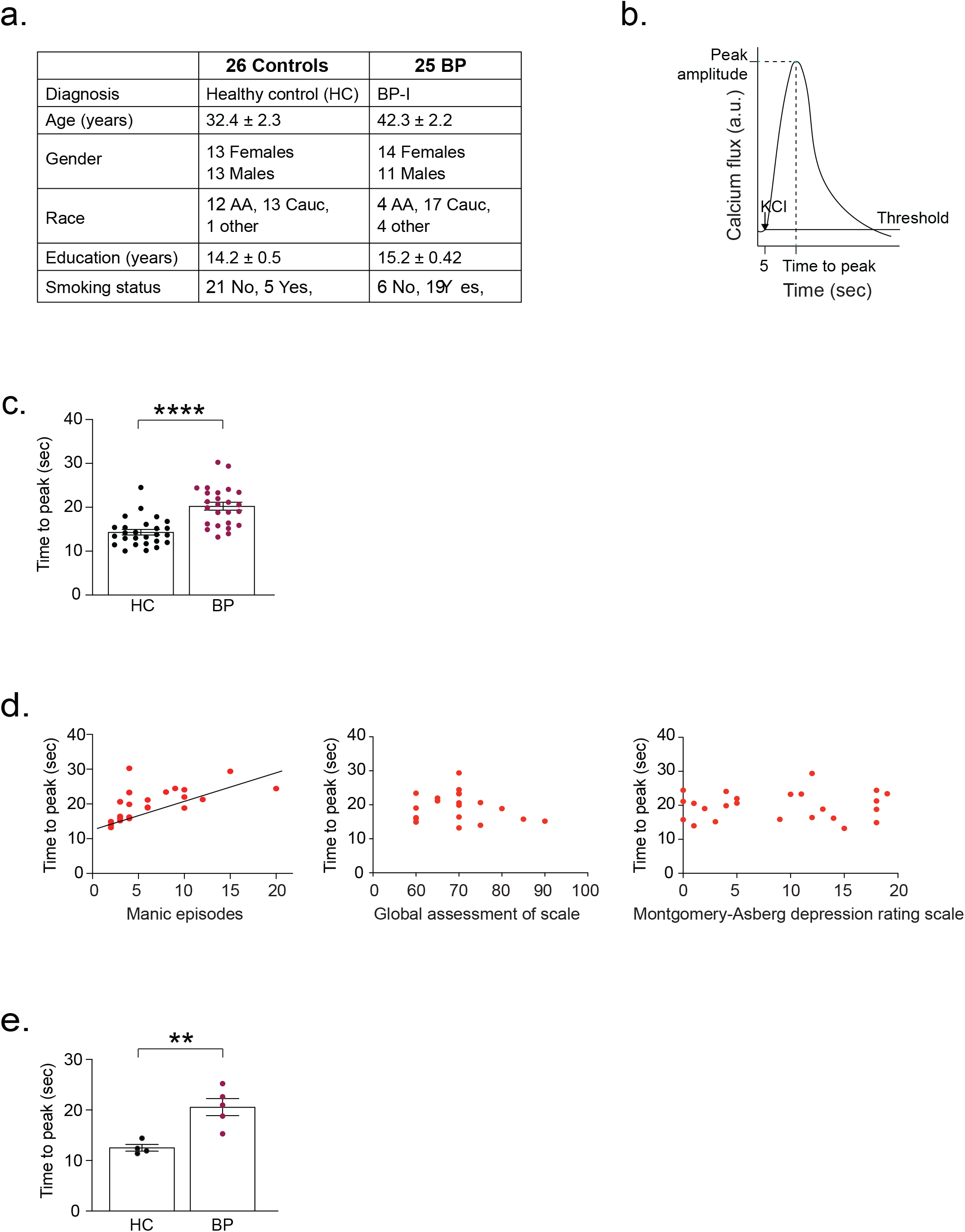
Deficit in the intracellular Ca^2+^ homeostasis in BP neurons. (**a**) Demographic distribution of the subjects included for the induced neuronal cells (iN cells) initial screening experiments; (**b**) Schematic depicting high potassium-evoked depolarization [Ca^2+^]_i_ homeostasis. Time to peak is measured for the first peak evoked; (**c**) BP derive excitatory iN cells showed delayed depolarization-induced time to peak. Each dot represents average for each subject. HC: Healthy Control; BP: Bipolar disorder-1; (**d**) correlation analysis between cellular phenotype and clinical phenotype showed significance between number of mania and time to peak after adjustment of several factors; (**e**) iPSC lines were generated for selected BP with best fit for correlation (d), and matched HC. Excitatory iPSC-neurons from BP again showed delayed depolarization-evoked [Ca^2+^]_i_ homeostasis.

### Attenuated CaV1.2-BK protein interactions in BP neurons

We hypothesized that this [Ca^2+^]_i_ deficit may be elicited by aberrant protein interactions involving CaV1.2. To address this question, we conducted a bioinformatics search using STRING database and found that CaV1.2 protein interacted with no more than 20 proteins with the highest confidence score of 0.9. Among these, we discovered one protein that has been known to regulate [Ca^2+^]_i_, which is the calcium and voltage activated potassium (BK) channel subunit KCNMA1^**6,7**^ (**Fig. 2a**).

**Fig. 2:**
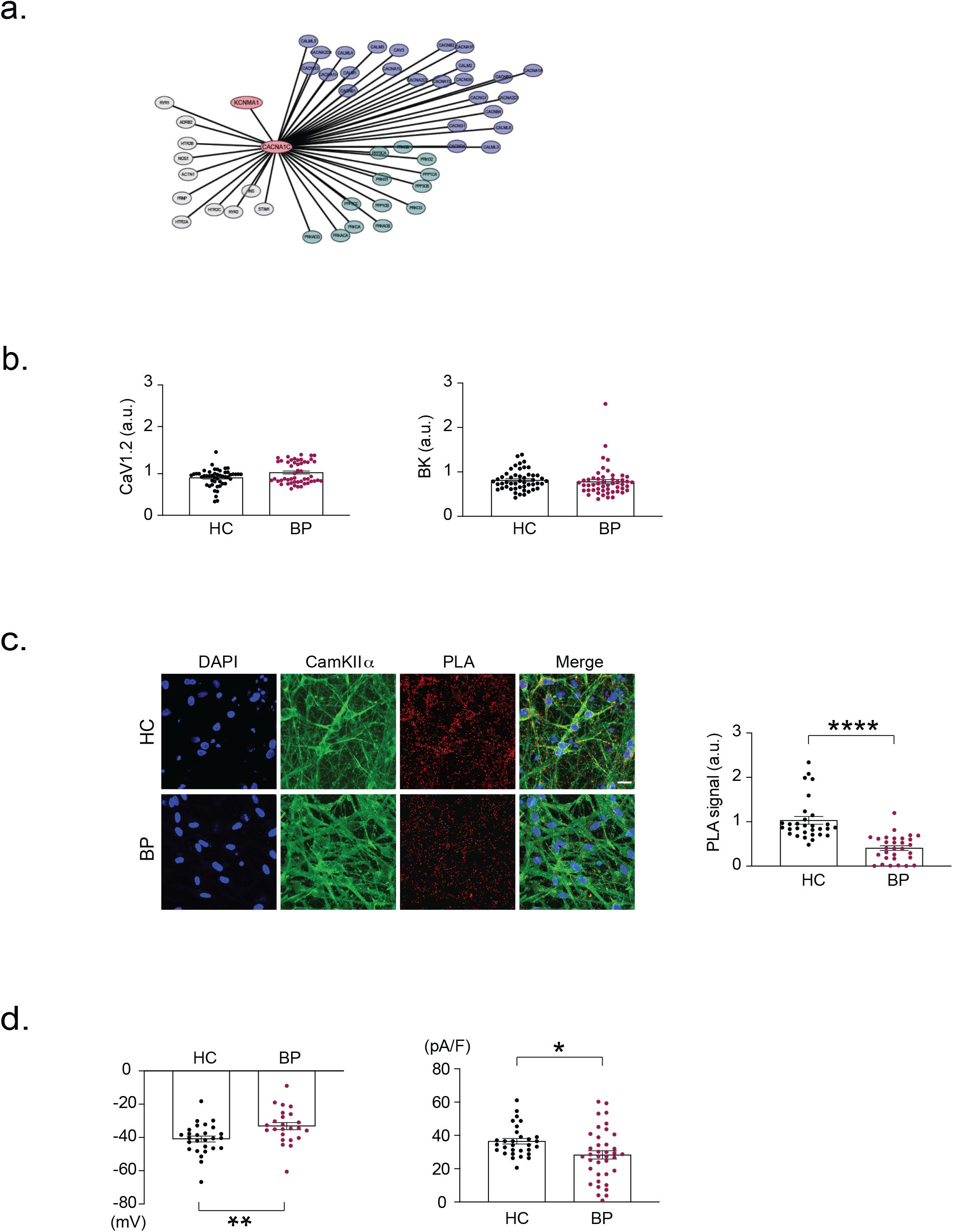
Attenuated CaV1.2-BK protein interactions in BP neurons. (**a**) STRING database hits for the CACNA1C (CaV1.2) interactors highlighted KCNMA1 (coding for the pore-forming subunit of the big potassium (BK) channel; (**b**) Immunofluorescence comparison for the CaV1.2 and BK channel for HC versus BP excitatory iPSC-neurons shows no significant difference in average image intensity normalized to CaMKII_α_. Similar results were obtained by western blotting (data not shown); (**c**) *In situ* proximity ligation assay (PLA) marking the interaction between CaV1.2 and BK channel as the red puncta dots shows significant decrease in interaction for the BP condition. Each dot here represents one image average; (**d**) Whole cell patch clamp electrophysiology showed a more depolarized state for the BP neurons and also decreased outward currents. Each dot represents neuron.

Accordingly, we tested whether CaV1.2-BK protein interaction may be altered in BP by using *in situ* proximity ligation assay (PLA). In iPS cell-derived excitatory neurons, we did not observe difference in the expression levels of these two proteins, respectively (**Fig. 2b**). However, the PLA signals indicated that the protein interaction was attenuated (**Fig. 2c**). The levels of the protein interaction were correlated with the number of manic episodes in BP patients (data not shown). Basic physiology has shown that BK channel outward potassium current is reduced when this protein is attenuated^**7**^. To test whether this reduction may also occur in BP neurons in which the pathological attenuation of CaV1.2-BK protein interaction is observed, we conducted whole cell patch clamp electrophysiology in BP patient neurons. We found that BP neurons were indeed more depolarized and had lesser outward potassium currents than the HC neurons (**Fig. 2d**), which are consistent with the theory established in basic physiology. Altogether, we demonstrate that CaV1.2-BK protein interaction is attenuated at both molecular and functional levels in BP patients.

### Attenuated CaV1.2-BK protein interactions as a driver for BP-associated intracellular Ca^2+^ homeostasis deficit: amelioration by BK-associated pharmacological interventions

We next wished to test whether the attenuated protein interaction is an upstream driver of delay of [Ca^2+^]_i_ time-to-peak in a high potassium-evoked depolarization, which is seen in BP patients in association with the number of manic episodes. To address this question, we aimed to enhance CaV1.2-BK protein interaction by introducing a linker protein between these two. The linker includes BK channel interactor (peptide from the auxiliary protein beta1)^**8,9**^ and CaV1.2 channel interacting nanobody^**10**^. Indeed, the introduction of this linker has enhanced CaV1.2-BK protein interaction evident by the enhanced PLA signals (**Fig. 3a**). Most importantly, this molecular intervention could ameliorate the delayed time-to-peak in BP patients, fixing this characteristic to the level seen in heathy subjects (**Fig. 3a**). These results clearly indicate that the attenuated CaV1.2-BK protein interaction is a key driver for the BP-associated intracellular Ca^2+^ homeostasis deficit.

**Fig. 3:**
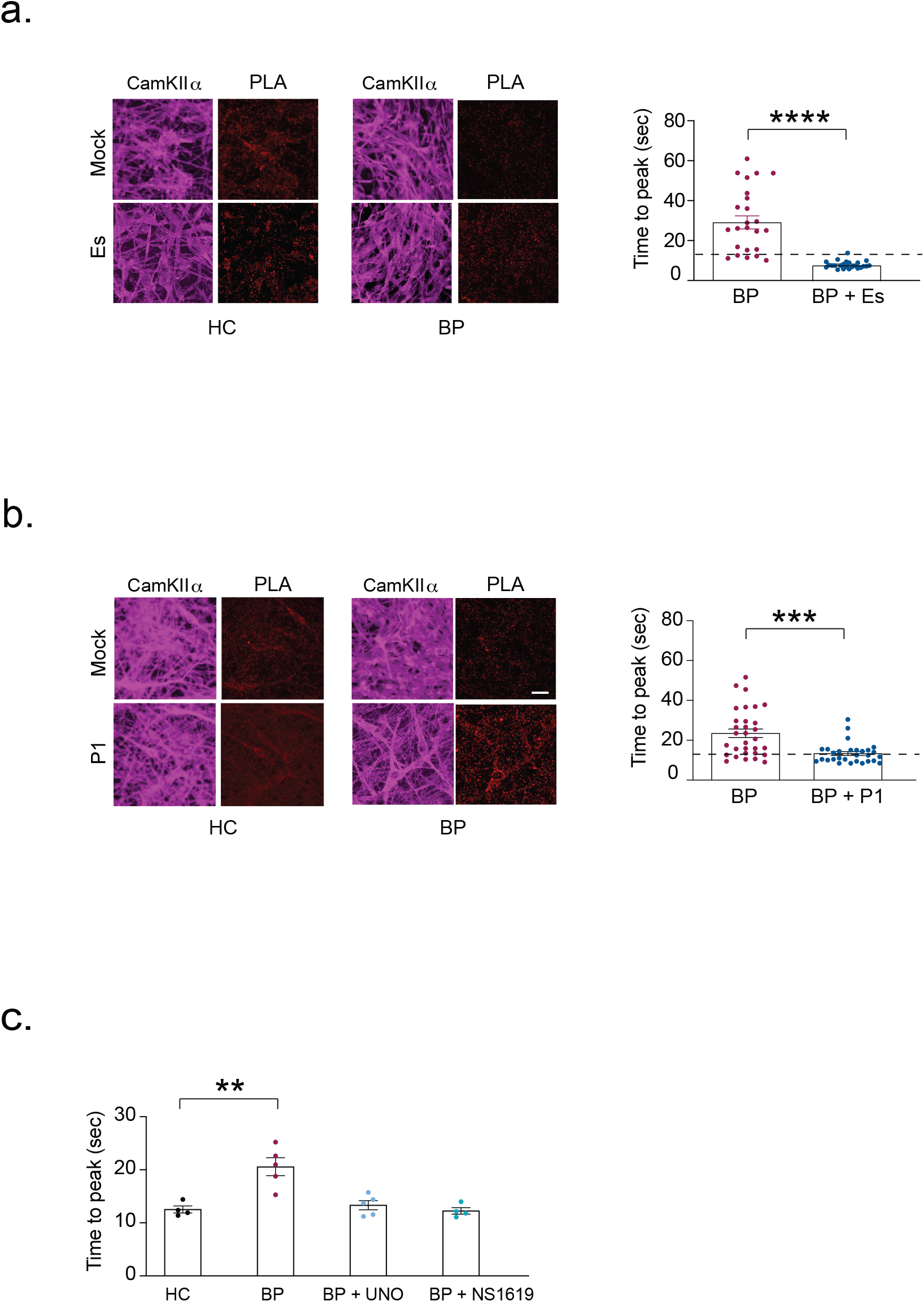
Attenuated CaV1.2-BK protein interactions as a driver for BP-associated intracellular Ca^2+^ homeostasis deficit: amelioration by BK-associated pharmacological interventions. (**a**) Pharmacological intervention with estramustine (Es), our bioinformatics-based hit that may affect the interaction, showed increased interaction and a resulting rescue of time to peak delay in two BP subject neurons. Dotted line represents HC average; (**b**) Molecular re-instatement of CaV1.2-BK channel interaction in BP iPSC-neurons by using a hybrid protein beta1-nanobody (P1). This also resulted in a rescue of time to peak phenotype as depicted for one BP subject. Scale bar is 20 μm; (**c**) Pharmacological interventions directly enhancing BK channel activity, viz. Unoprostone (Uno) and NS1619, also showed a rescue effect on time to peak in BP subjects. Each dot represents here average of one BP subject.

Encouraged by this data that the enhancement of the attenuated CaV1.2-BK protein interaction is a way of normalizing the key cellular deficit that may be associated with clinical cycling features in BP, we looked for compounds that may have a similar action. To address this, we looked for compounds that interest with BK channel. We conducted a bioinformatic assessment using BioGRID4.4 database and underscored estramustine and fluticasone propionate. In particular, as a cytoskeletal anchor that interacts with the BK channel subunit KCNMA1, we hypothesize that estramustine may affect CaV1.2-BK protein interaction. At the experimental level, we observed that the BP-associated attenuation of the protein interaction was ameliorated (**Fig. 3b**).

Importantly, once the protein interaction is normalized by estramustine, we also observed amelioration of the disease-associated [Ca^2+^]_i_ deficit (**Fig. 3b**).

We also tested whether other interventions targeting to BK channel may also be effective to ameliorate the disease-associated [Ca^2+^]_i_ deficit. To address this question, we used two types of BK channel activators unoprostone isopropyl ester (Uno) or NS1619. First, the addition of BK channel activators ameliorated the aberrant BK-associated potassium current characteristics that exist in associated with the attenuated CaV1.2-BK protein interaction (data not shown).

Importantly, we also observed that both Uno and NS1619 were effective to ameliorate the [Ca^2+^]_i_ deficit in BP neurons (**Fig. 3c**).

### Stress-induced behavior phase shift in mice: amelioration by BK-associated pharmacological interventions

Thus far, we have demonstrated the [Ca^2+^]_i_ homeostasis-related cellular deficit that is clinically associated with manic episode numbers. For this cellular deficit, we underscore disease-associated attenuation of CaV1.2-BK protein interactions as a key upstream driver. This [Ca^2+^]_i_ deficit can be ameliorated by targeting to CaV1.2-BK protein interaction or more directly to the BK channel. Altogether, our cellular observations linking to the clinical phenotypes highlight BK channel potentially as a key molecule for mood cycling. Thus, we aimed to model this cycling in mice by using BK channel as a lead molecule. Clinically, mood cycling frequently occurs in association with stressors. Altogether, we combined KCNMA1(BK) knockout mice with a mild stressor of a short sleep deprivation (SD) possibly to model cycling (**Fig. 4a**).

**Fig. 4:**
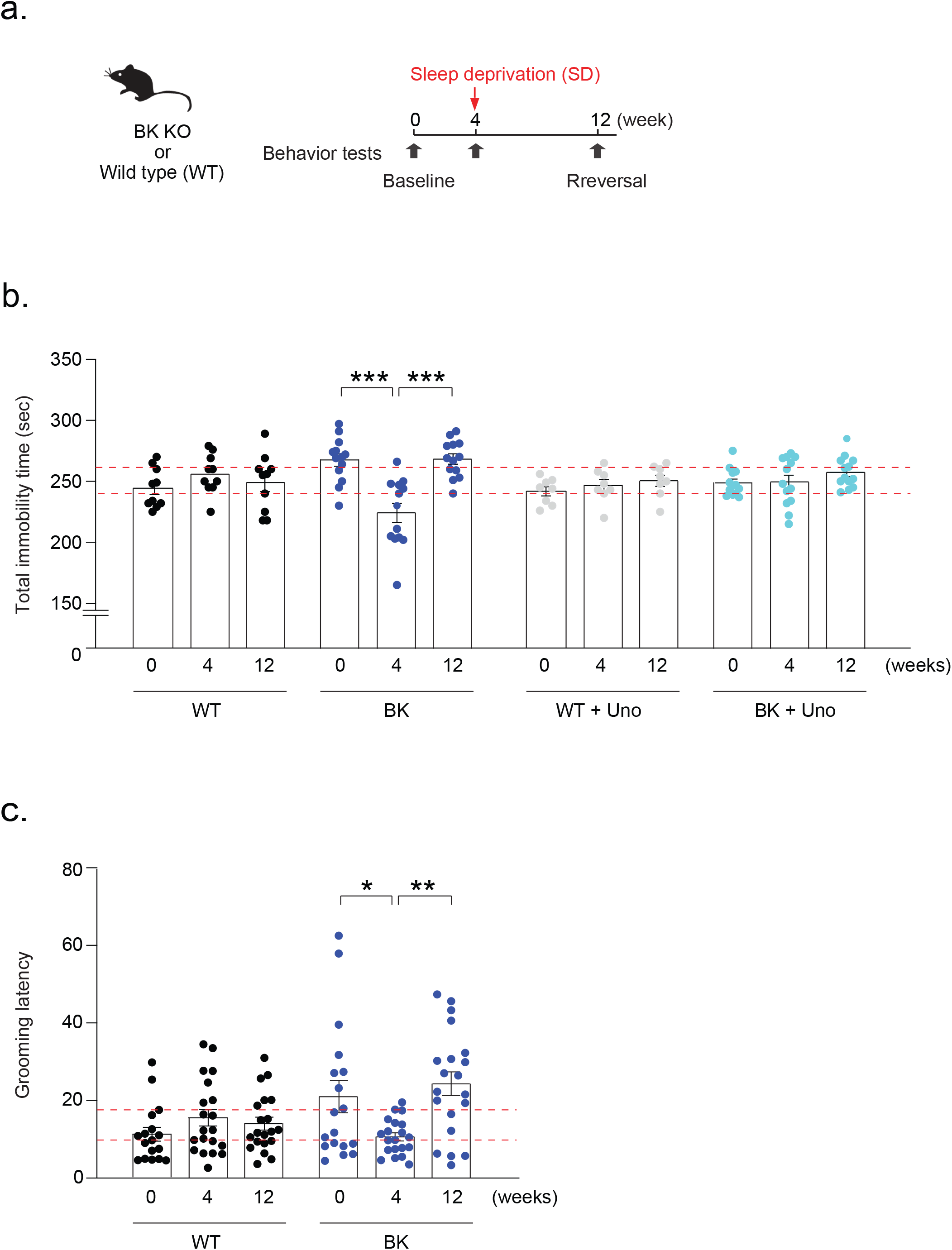
Stress-induced behavior phase shift in mice: amelioration by BK-associated pharmacological interventions. (**a**) Behavior phase switch study paradigm schema. The mice are first tested at baseline behavior in week 0, followed by acute sleep deprivation and re-testing in week 4. This is followed by a long rest and re-testing in week 8; (**b**) CaMKII_α_-driven *KCNMA1* knockout heterozygous (BK) mice were tested for the behavior phase switch with forced swim test and showed a ‘phase switch’ with sleep deprivation of 4 hours. Unoprostone (Uno) was able to rescue this switch in knockout mice. Notably, no significant difference in behavior was observed for the wild type (WT) mice; (**c**) Behavior phase switch was also observed when the BK mice performed sucrose splash test. Each dot represents one mouse.

We observed that heterozygous knockout of BK mice (BK Het) showed significantly higher immobility compared with control mice in forced swim test at the baseline (**Fig. 4b**). Four weeks after we examined the baseline behaviors, we applied acute SD for four h to both BK Het and WT, respectively. This mild stressor did not change the behavior in WT when we monitored this immediately after SD. In contrast, we observed a significant change in FST behaviors in BK Het: although more immobile compared with WT at the baseline as described above, BK Het became less immobile compared with WT immediately after SD (**Fig. 4b**). This change looks like a “phase switch” in BK Het compared with WT. This switch is unlikely a mere acute response, as the less immobile status stayed at least for one week as far as we tested (data not sown). Interestingly, this stress-induced status in BK Het disappeared and went back to the default status (baseline level) eight weeks after the SD-elicited switch. Indeed, without any further intervention after the SD, BK Het displayed significantly higher immobility compared with WT again (**Fig. 4b**). Similar behavioral phase switch was also observed in sucrose splash test (**Fig. 4c**).

Finally, we tested whether the pharmacological intervention with BK channel may ameliorate this “phase switch”. Thus, we intraperitoneally (I.P.) injected Uno (0.06 mg/kg) or saline 20 min before behavior testing at each stage (baseline, SD, and default). Interestingly, we found not only the drug was able to rescue the higher immobility phenotype at baseline and default stage (0 and 12 w, respectively: see **Fig. 4b**), but was also able to rescue the switch due to SD (4 w: see **Fig. 4b**). WT mice had no statistically significant effect with these injections (**Fig. 4b**).

### Novel mechanism for BP with an association with mood cycling: a possible amelioration by BK-associated pharmacological interventions

The recurrent cycling is known to be associated with poor prognosis. Thus, it is important to decipher the pathophysiological mechanism that may possibly underlie the cycling for a future treatment of BP. Here we report intracellular Ca^2+^ homeostasis deficit in BP patients, which is primarily driven by the attenuated CaV1.2-BK protein interaction. Intervention with the protein interaction or directly with BK channel can ameliorate the intracellular Ca^2+^ homeostasis. Given that the cellular deficit is associated with the number of manic episodes, suggesting the implication in mood cycling, we developed a novel mouse model of phase switch. Interestingly, this mouse behavior was also ameliorated by targeting to BK channel. Thus, further investigation of this molecular and cellular cascade may be a promising means for translational research for BP.

## ACKNOWLEDGEMENT

We thank Ms. Yukiko Y. Lema for manuscript organization. We also thank Dr. Fernando Goes, Dr. Hanna Jaaro-Peled, Dr. Kun Yang, Dr. Manu Ben-Johny, Dr. Anjali Rajadhyaksha, Dr. Andrea Meredith, Ms. Josephine de Chabot de Tramecourt, Ms. Lauren Guttman, Mr. Imad Isehak, as well as many other Sawa lab members and collaborators for scientific contributions. This observation has been included in the patent application filed as PCT/US2024/053139.

## METHODS

### Subject recruitment and assessment

Johns Hopkins Schizophrenia Center recruited control and BP patients for this study and all the protocols were approved by the Johns Hopkins Medicine Institutional Review Board (IRB). Prior written informed consent in each case was obtained from the participants after discussing the purpose, sample extraction procedures and all other details of the recruitment procedure. Trained dermatologists performed the skin biopsy as per approved protocol. A total of 26 control and 25 BP-type 1 patient samples were utilized in the study. All clinical scales were administered by board-certified psychiatrists. The diagnosis of BP was assessed by the Diagnostic Interview for Genetic Studies (DIGS) 4.0 interview; the presence and severity of depressive and manic symptoms was measured respectively by the Montgomery Asberg Depression Rating Scale (MADRS) and the Young Mania Rating Scale (YMRS). Finally, the Retrospective Criteria of Long-Term Treatment Response in Research Subjects with Bipolar Disorder (Alda scale) was used to evaluate treatment response to lithium.

### Antibody

The primary antibodies used for human neuronal cultured cells included: CaMKII_α_ (R&D systems, MAB5584), MAP2 (Sigma, M2320), GFAP (Dako, Z0334), GAD67 (Millipore, MAB5406), vGLUT1 (Abcam, ab72311), synapsin-1 (Millipore, AB1543), PSD-95 (ABcam, ab2723), BRN2 (Cell Signaling tech. 12137S), SATB2 (ABcam, ab51502), CTIP2 (ABcam, ab18465), Anti-CaV1.2 (Alomone labs, ACC-003-GP), and anti KCNMA1 (Alomone labs, APC-107) were used for the immunofluorescence, proximity ligation assay, co-IP studies.

### Plasmids, and virus

Plasmids for the study have been prepared with the PureYield^TM^ plasmid midiprep system (A2492) (Promega). Lentiviral particles harboring human *ASCL1, POU3F2* and *MYT1L* and were generated as previously described^**11**^ and were used at 1:4 multiplicity of infection (MOI). pLenti-CaMKIIa-GCaMP5g was generated by sub-cloning GCaMP5g from pCMV-GCaMP5G [(pCMV-GCaMP5G was a gift from Douglas Kim & Loren Looger & GENIE Project (Addgene plasmid # 31788; http://n2t.net/addgene:31788; RRID:Addgene_31788)]^**12**^ into pLenti-CaMKIIa-ChETA-EYFP [pLenti-CaMKIIa-ChETA-EYFP was a gift from Karl Deisseroth (Addgene plasmid # 26967; http://n2t.net/addgene:26967; RRID:Addgene_26967 (Addgene plasmid # 26967)]^**13**^. It was used at 1:8 MOI as determined by the functionality in the neuronal cells. pLenti-CaMKIIa-mCherry was generated by sub-cloning mCherry from ChR2-mCherry-rAAV vector^**14**^ into FCK(1.3)GW lentivector^**15**^. Beta1-SH3 construct generated with Twist Biosciences (https://www.twistbioscience.com/) using pTWIST-CMV-beta globin backbone.

### Immunoprecipitation and western blotting

Cells were lysed in lysis buffer (150 mM NaCl, 50 mM Tris-HCl, pH 7.5, 1% Triton X-100, will be modified) containing a protease inhibitor mixture (Roche Applied Sciences). Lysates were sonicated, cell debris was cleared by centrifugation, and the soluble fraction was immunoprecipitated as described previously^**16**^.

### Fibroblast generation and cell culture

Skin biopsies were done by the trained clinician staff at the Johns Hopkins, and the fibroblast cell lines were generated with the Johns Hopkins core. The fibroblast lines were maintained with MEM backbone media supplemented with 10% FBS, 1X MEM-NEAA, 1X pen-strep and 1X sodium pyruvate.

### Coverslip preparation

iN cell and iPSC-neuron differentiation was performed on 12 mm glass coverslips coated with 100 µg/ml poly-L-ornithine and 5 µg/ml laminin (PO/L) Other cell culture surfaces (like, six-well plate or culture flask etc.), were similarly coated with PO/L when stated.

### Fibroblast derived iN cells

iN cells were generated with some modification to our published protocol^**11**^. 20,000 fibroblast cells were transduced with *ASCL1, POU3F2, MYT1L* (1:4 MOI). Small molecule (SMC) media comprised of Neurobasal medium (50% backbone), DMEM-F12 (50% backbone), N2 supplement (1X), B27 supplement (1X) glutaMAX (0.5X), ITS-X (0.5X), db-cAMP (200 µM), CHIR99021 (2 µM), SB431542 (10 µM), Noggin (500 ng/ml) and lamimin (1 µg/ml). SMC was maintained for two weeks. The final (FD) media was then started and maintained for additional four weeks, and comprised of Neurobasal (100% backbone), B27 supplement (1X), GlutaMAX (1X), BDNF (10 ng/ml), GDNF (2 ng/ml), NT3 (10 ng/ml)and laminin (1 µg/ml).

### iPSC generation and cell culture

The skin fibroblasts from BP patients at early passage (p2) were used for the generation of iPSC. The ployclonal iPSC lines were generated by the New York Stem Cell Foundation (NYSCF) according to their published protocol^**17**^.

### iPSC derived excitatory neuron-enriched culture generation

STEMdiff neural system (StemCell technologies) with minor modifications was used to generate central nervous system-type neural progenitor cells (NPC). Briefly, iPSC lines were seeded on PO/L coated wells of the six-well plate in the presence of STEMdiff neural induction medium + SMADi (twice). The cells were re-passaged and cultured in STEMdiff forebrain neuron differentiation media until confluency. These NPC could be further passaged and cryopreserved with Synth-a-Freeze until further differentiation. Thiazovivin (1µM) was included in media each time while passaging or thawing the cells. The media composition used for FD stage included BrainPhys backbone (100%), glutaMAX (1X), B27 supplement (1X), BDNF (10 ng/ml), GDNF (2 ng/ml), NT3 (10 ng/ml), laminin (1 µg/ml) as adapted from the iN FD media to generate CaMKII_α_ positive neurons, and additionally Ascorbic acid and db-cAMP.

### Proximity ligation assay (PLA)

iPSC-neurons were fixed with 4% paraformaldehyde (PFA) and PLA was performed with Sigma’s Duolink *in situ* PLA reagents. The signal visible as the distinct fluorescent spot was visualized with LSM700 confocal microscope at 40X magnification and quantified either for average image intensity by using Fiji software (imagej.net) or puncta count done by software Volocity.

### Calcium imaging

Olympus fixed-stage BX61-Wi microscope with infrared-sensitive CCD camera (iXon3, ANDOR technology) set-up on anti-vibration table, and high speed galvo mirror with filters (DG4-1015, Shutter Instruments Company) were used for all the imaging. CaMKII_α_-driven genetically encoded calcium indicator version 5g (GCaMP5g) has been used as the Ca^2+^ change marker in the neuronal cells for high potassium-mediated depolarization-evoked Ca^2+^ kinetics. Normal artificial cerebro-spinal-fluid (aCSF) solution composed of NaCl (150 mM), KCl (3 mM), CaCl2.2H_2_O (2mM), MgCl2.6H_2_O (1.3 mM), HEPES (10 mM) and glucose (10 mM).

High potassium solution was composed of NaCl (53 mM), KCl (100 mM), CaCl2.2H_2_O (2mM), MgCl2.6H_2_O (1.3 mM), HEPES (10 mM) and glucose (10 mM). Osmolarity was set between 290 to 300 mOsmol/kg for all the solutions (Advanced instruments) and pH set to 7.4 (Fisher) with NaOH. 1 µM tetrodotoxin was maintained in all the solutions for calcium imaging. 401 images were taken for calcium kinetics. Images were processed through the Metamorph advanced, Molecular device. For kinetics dF/F_base_ values were calculated for the region of interest selected in the cell body near extension junction. Only the cells having dF/F_base_ values above the threshold cutoff of 5% were used, and only the first peak values were included for the analysis. Cells transduced with lentivirus harboring CaMKII_α_-mCherry were used for the baseline [Ca^2+^]_i_ measurements. For loading, Fura2-AM 1 mM stock was made in DMSO anhydrous, and 1 μM working was made in FD media. Imaging was completed within 1 hour of dye loading. All drugs were used acutely 30 min before all the recordings and maintained in all the solutions. Drug doses Uno (2 nM), NS1619 (10 μM), NS309 (5 μM), dantrolene (50 μM), thapsigargin (10 μM), 2-APB (10 or 100 μM as specified), verapamil (10 μM), ω-agatoxin TK (1 μM), huwentoxin XVI (150 nM), SNX-482 (100 nM), FPL64176 (1 μM), lithium (1 mM), carbamazapine (100 μM), phenytoin (100 μM), VPA (500 μM), iberiotoxin (100 nM), estramustine (10 μM).

### Electrophysiology

Coverslips containing iN cells or iPSC-neurons were loaded on an upright microscope (Olympus BX-50-WI) fitted with a Warner RC-27 submerged recording chamber. Solutions and recording conditions were adapted from the published article^**18**^. Patch pipettes (5-8 M_Ω_) were pulled and filled with an internal solution composed of potassium gluconate (126 mM), KCl (8 mM), HEPES (20 mM), EGTA (0.2 mM), NaCl (2 mM), MgATP (3 mM), and Na_3_GTP (0.5 mM) (pH 7.3, 290-300 mOsm). Recordings were made at 23-25°C in aCSF composed of NaCl 137 mM, KCl 5 mM, CaCl_2_ 2 mM, MgCl_2_ 1 mM, HEPES 10 mM, and glucose 10 mM (pH 7.4, 290-300 mOsmol/kg). CaMKII_α_-promoter driven mCherry positive neurons were identified by epifluorescence signal through CCD camera (ANDOR-iXon3). Whole-cell patch-clamp recording were obtained with a Multiclamp 700B amplifier (Molecular Devices) controlled by pClampex 10.3 (Axon Instruments), and acquired at a sampling rate of 10 kHz with Digidata 1440A (Molecular Devices). All recordings were made in the presence of TTX (1 μM) unless otherwise specified. Cells were held at approximately −70 mV, then a series of voltages steps from −100 to +60 mV (in 20 mV increments for 500 ms) were delivered through the patch electrode for current recording. The peak amplitude of the resulting outward currents was quantified with leak subtraction and normalized with cell capacitance.

### Mice

Animals were group housed after weaning and maintained on a 12-hour light/dark cycle with food and water provided *ad libitum*. All mice were used in accordance with Johns Hopkins University School of Medicine Institutional Animal Care and Use Committee (IACUC) guidelines. Mice testing was performed in batch of 8-15 mice, with the earliest starting age around 12 weeks.

Kcnma1(fl/fl)-tdTomato (fBK) mice were provided by their generator, Dr. Andrea Meredith, University of Maryland^**19**^. Genotyping was performed with N1: 5_′□_CAAAGGGGGTTTGCTTGTGAGAGG_□_3_′_ and N2:

5_′□_CATGAGCGTGTGCCTAAACGCA_□_3_′_, which amplify a 642□bp product from the Kcnma1-tdTomato floxed conditional allele and a 577□bp product from WT controls.

Mice for comparing WT, Het (fl/+) and Hom (fl/fl) mice were generated by fl/+ x fl/+ matings for injection of a control virus [pENN.AAV.CamKII0.4.eGFP.WPRE.rBG was a gift from James

M. Wilson (Addgene plasmid # 105541; http://n2t.net/addgene:105541; RRID:Addgene_105541)] or a CaMKIIa-Cre virus [pENN.AAV.CamKII.HI.GFP-

Cre.WPRE.SV40 was a gift from James M. Wilson (Addgene viral prep # 105551-AAV9; http://n2t.net/addgene:105551; RRID:Addgene_105551]. For generation of CaMKIIa-Cre x fBK mice we crossed Hom B6.Cg-Tg (CaMKIIa-cre)T29-1Stl/J mice (Jax line 005359) with fl/+ BK mice and compared the resulting WT vs. f/+ mice (all were Cre/+).

Ear tagging was performed to achieve individual identification followed by utilization of the minute ear piece extracted during tagging for genotyping.

For mouse Genotyping, tissue samples were lysed into crude DNA using a PCR lysis reagent (Viagen: 102-T). Subsequently, crude DNA was diluted ten times and used in PCR protocols with Taq polymerase (Takara: RR001A) as necessary for either Cre or floxed genes.

### Behavior assays

#### Forced swim test (FST)

The FST was used to determine mice mobility as a proxy for anxiety-like behavior. Water was prepared overnight in 5000 ml beakers, animals were gently placed in the water and their activity was recorded with a camcorder for six minutes. Multistopwatch (http://www.multistopwatch.com/) was used for scoring the active struggling time.

#### FST and behavior phase switch paradigm

Throughout testing period, mice were provided food pellets and hydrogel *ad libitum*. For behavior phase switch, animals were exposed to repeated FST with stated interval between each trial. The test was started by measuring the baseline FST behavior in week 0 at ZT 4. In week 4, sleep deprivation paradigm was provided as a mild stressor before FST from ZT 0 (lights on time) to ZT 4, after which they were tested for FST. The phase of baseline-sleep deprivation-reversal testing was repeated after a period of 3 months where mentioned. Mice were given Uno (0.06 mg/kg) by intraperitoneal (IP) injection 20 minutes before behavior testing, where mentioned.

#### Sucrose Splash Test (SST)

Mice were first transferred out of their home cage and kept in a surrogate cage for 1 hour of acclimation. Home cages were then set up underneath a camera for behavior recording. Each mouse was sprayed on the back twice with a 10% sucrose solution made in distilled water. They were then placed gently in their home cages and recorded for 5 minutes. The behavior phase switch paradigm was repeated with this behavioral assay like for the FST. Grooming latency was recorded by using spacebarclicker.org.

### Data analysis

#### Ca^2+^ pharmacology and behavior analysis

Data analysis for Ca^2+^ pharmacology was performed in Graph pad prism with ANCOVA for multiple comparisons; and t-test analysis for in-group comparison and a p value of 0.05 was considered significant.

#### Correlation analysis of time-to-peak with clinical phenotypes

Linear regression analysis adjusting for age, gender, race, smoking status, duration of illness, cuprizone, and mood stabilizer usage (Yes/No) were conducted to test the correlations between time to peak and main clinical variables, including number of mania episodes, number of depressive episodes, number of total episodes, global assessment scale (GAS), young mania scale total, ALDA total score.

